## Supplementary Material for "Structure of the yeast Nup84-Nup133 complex details flexibility and reveals universal conservation of the membrane anchoring ALPS motif"

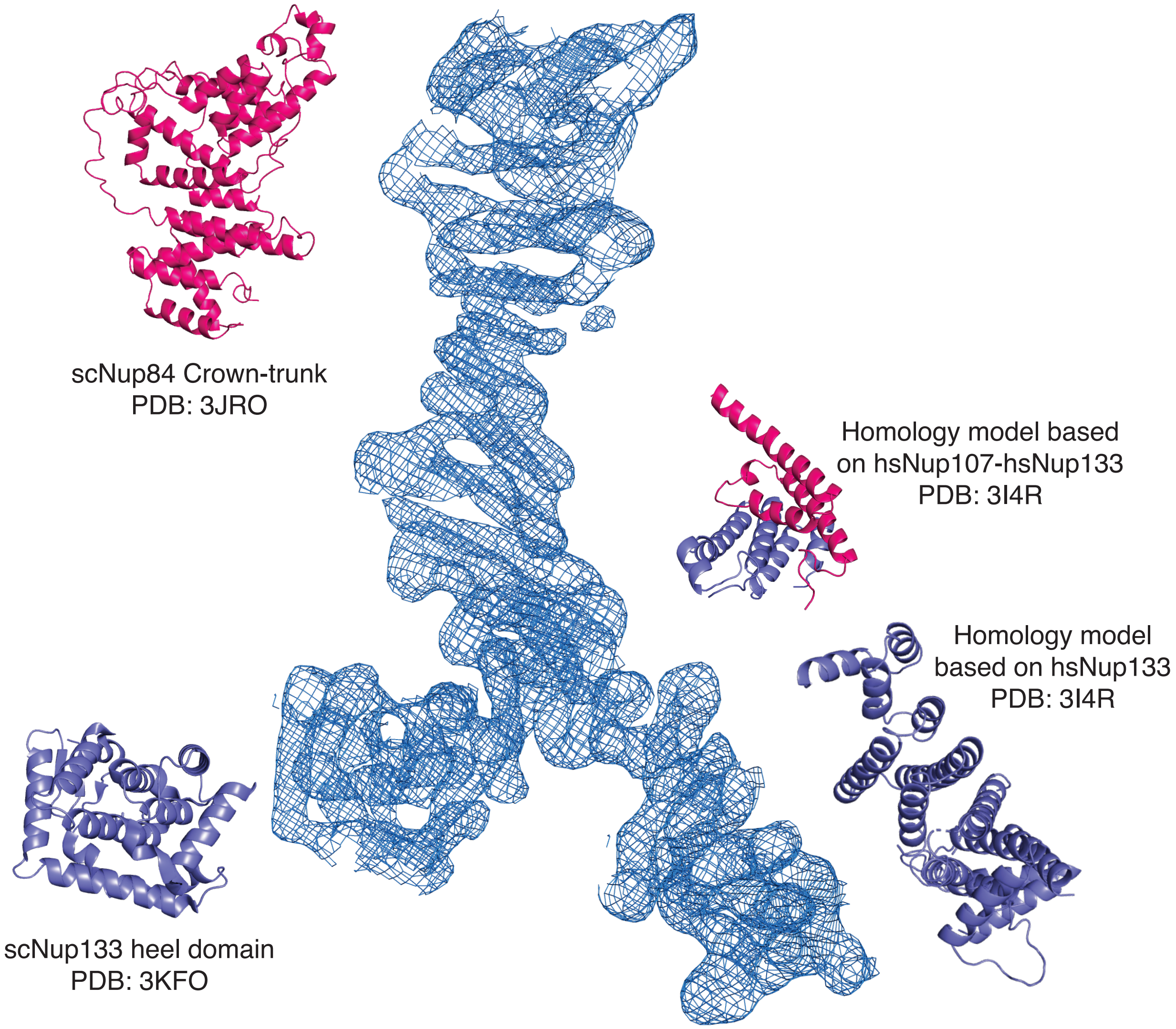


### Supplementary Figure 1.

**Electron density map for the Nup84-Nup133$_{CTD}$ structure.**
Final $2F_O$-$F_C$ electron density map contoured at 1.5 $\sigma$ with fragments used to build the structure. Density for VHH-SAN8 was excluded for clarity.

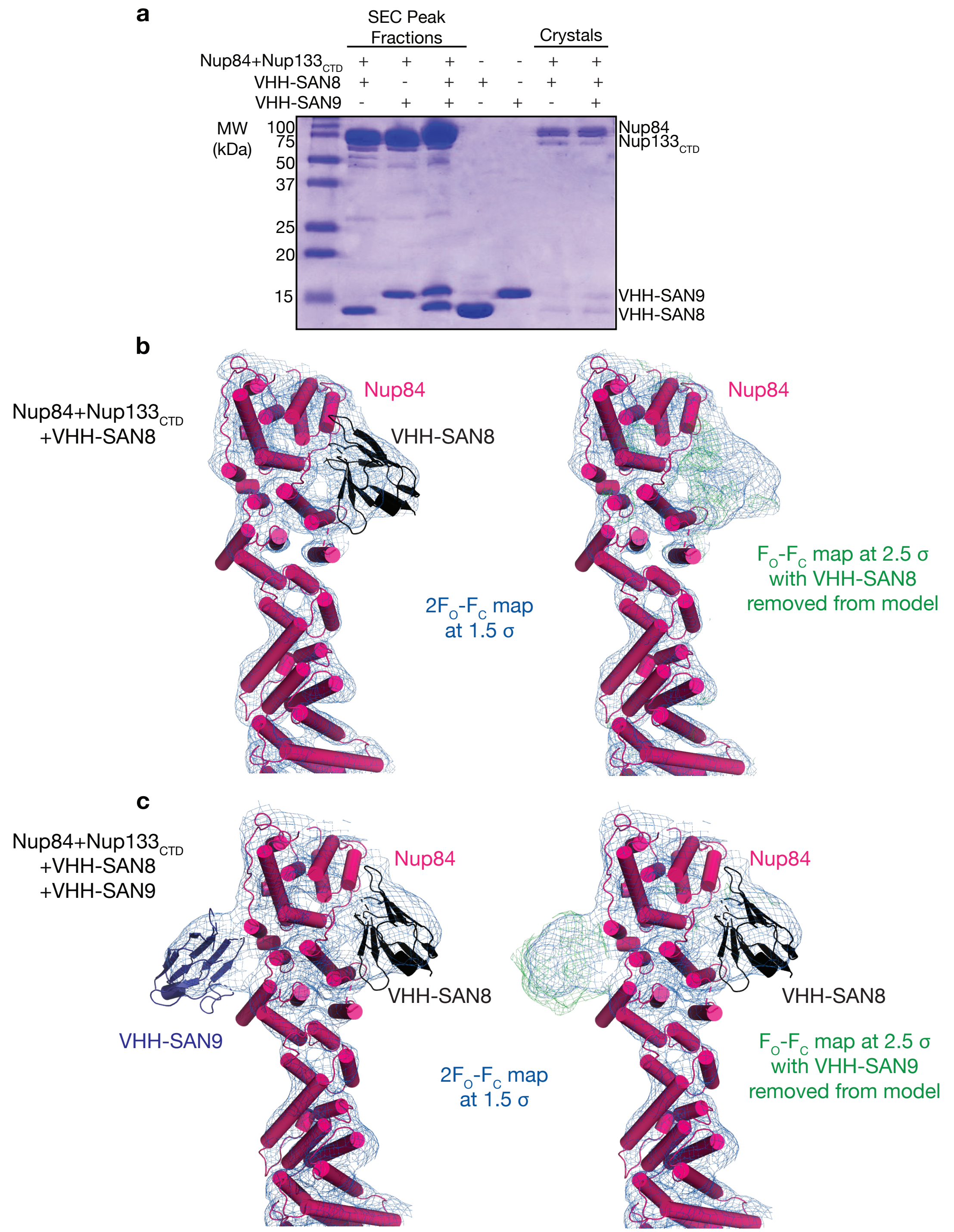


### Supplementary Figure 2.

**VHH-SAN8 and VHH-SAN9 are present in the structures.**

(a) SDS-PAGE gel of purified Nup84-Nup133$_{CTD}$-VHH complexes and crystals of Nup84-Nup133$_{CTD}$-VHH-SAN8 and Nup84-Nup133$_{CTD}$-VHH-SAN8/9 complexes. (b) Binding location of VHH-SAN8 shown by density in both 2F$_O$-F$_C$ (left) and F$_O$-F$_C$ omit map when VHH-SAN8 is removed from the structure (right). (c) Same analysis as in (b) with the structure of Nup84-Nup133$_{CTD}$-VHH-SAN8/9.

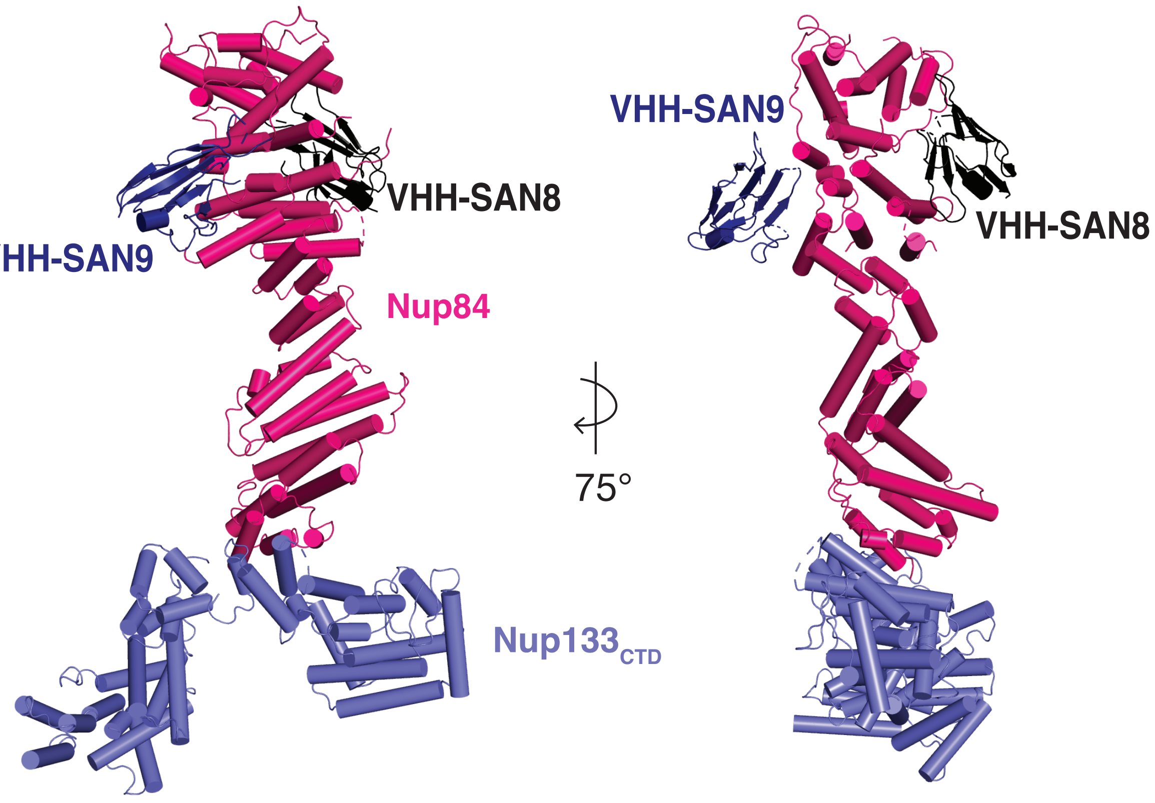


**Supplementary Figure 3.**

**Structure of the *S. cerevisiae* Nup84-Nup133$_{CTD}$-VHH-SAN8/9 complex.**

Structure of Nup84-Nup133$_{CTD}$-VHH-SAN8, with Nup84 in pink, Nup133$_{CTD}$ in purple, and VHH-SAN8 in black, VHH-SAN9 shown in navy.

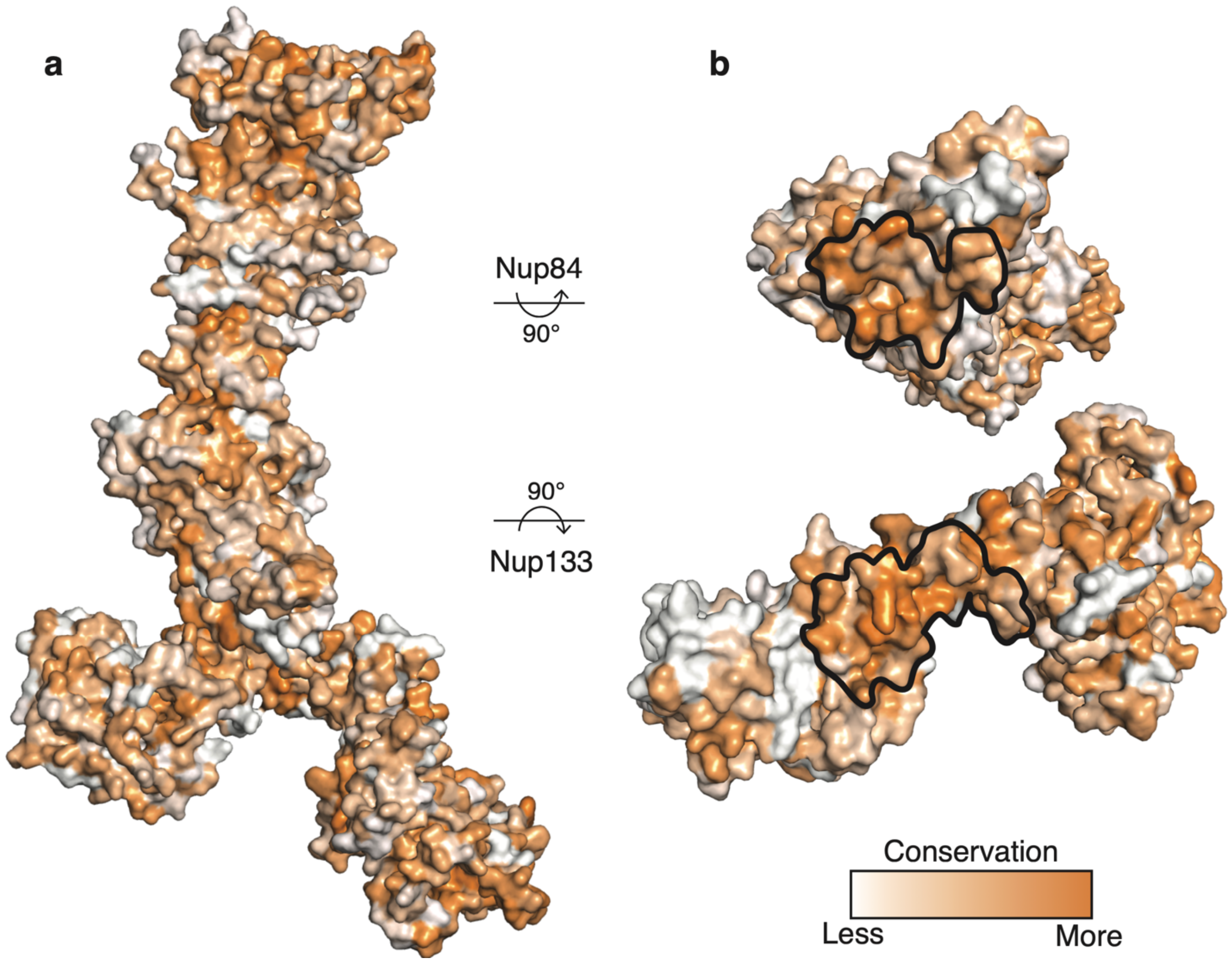


### Supplementary Figure 4.

**Conservation of Nup84-Nup133$_{CTD}$.**

(a) Surface rendering of Nup84-Nup133$_{CTD}$ with a color gradient depicting conservation across diverse eukaryotes. (b) Open-book depiction of the interface between Nup84 and Nup133. The binding interface is outlined in black.

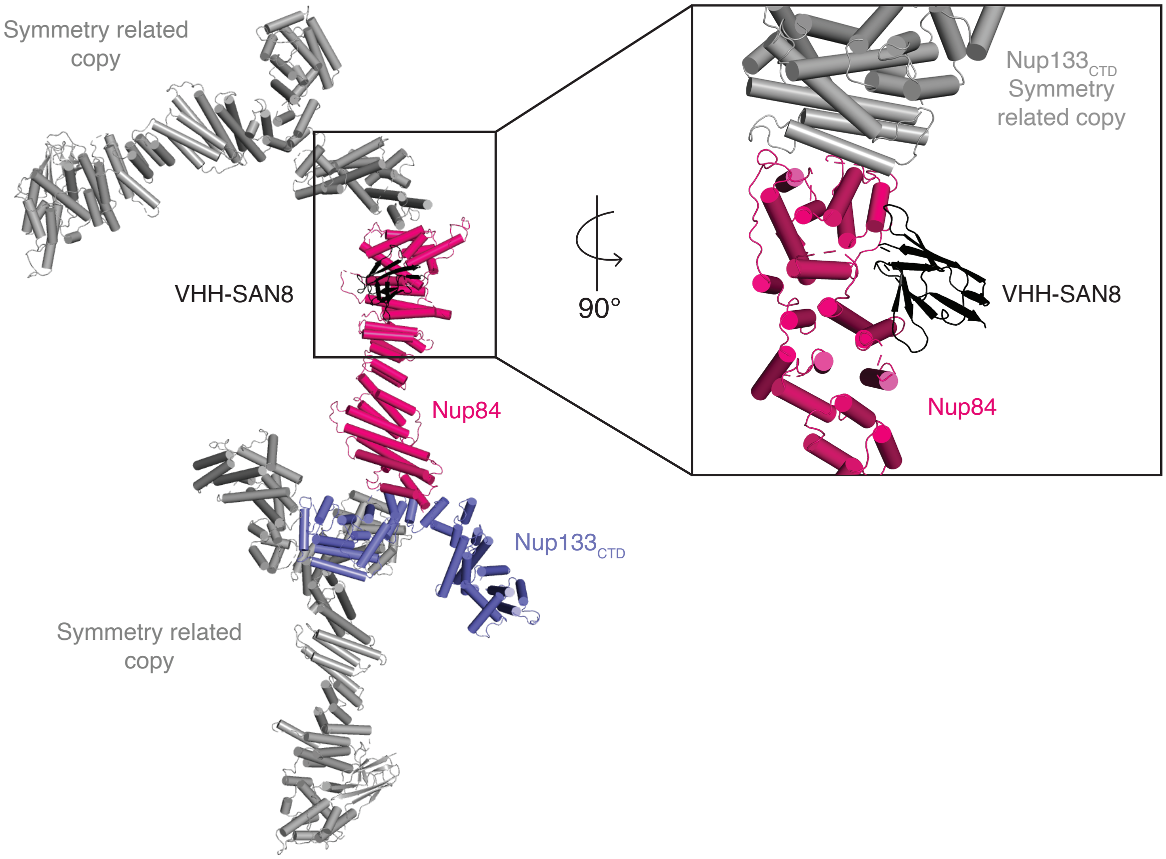


### Supplementary Figure 5.

**VHH-SAN8 is critical for rigidifying a crystal packing interface.**
Depiction of packing interactions within the crystal. Two symmetry related copies are shown for Nup84-Nup133$_{CTD}$-VHH-SAN8 (gray). Inset shows a closer view of the interaction between Nup84 and a symmetry related copy of Nup133$_{CTD}$.

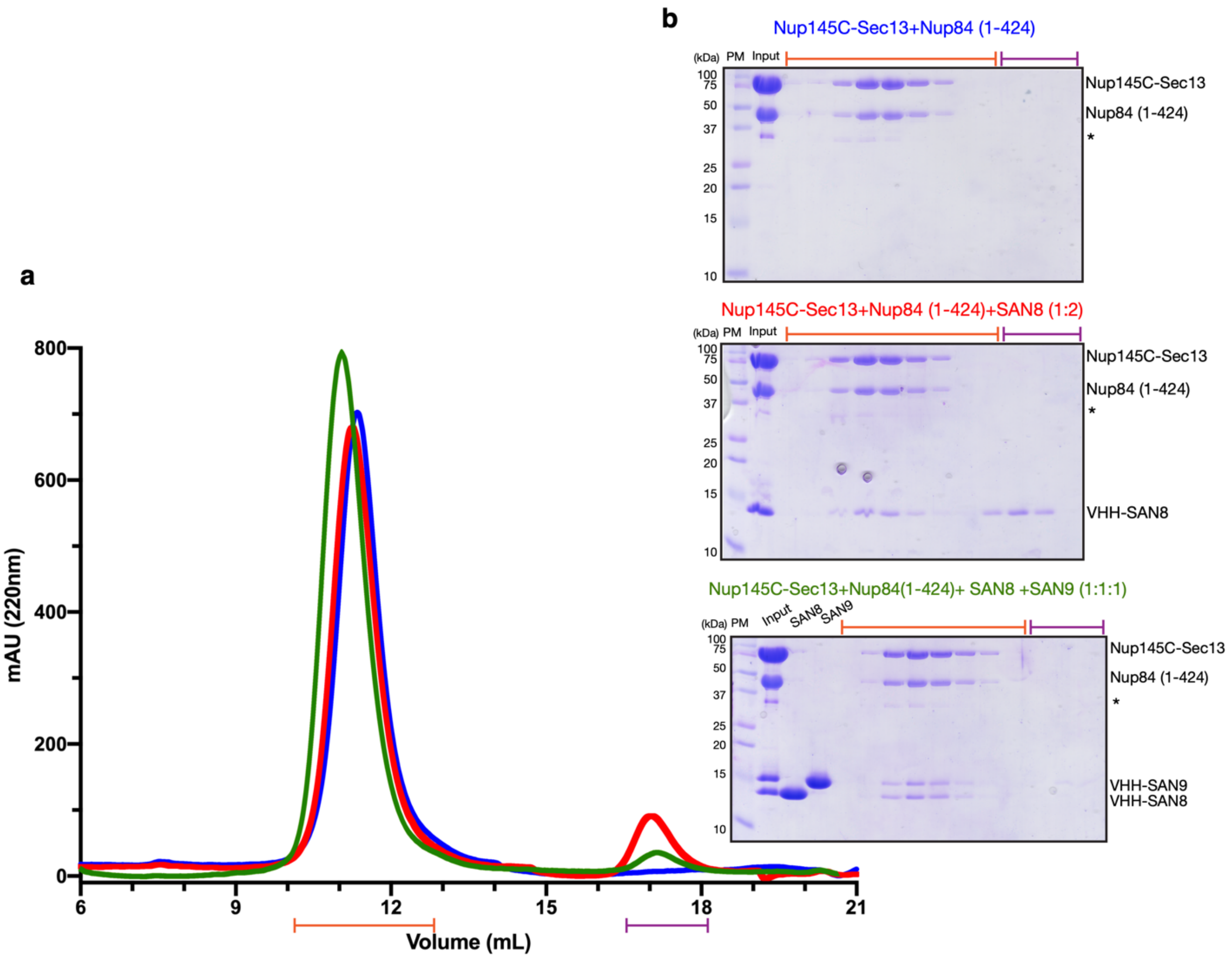


### Supplementary Figure 6.

**VHH-SAN8, VHH-SAN9 bind Nup84 with in complex with Nup145C.**

(a) Size exclusion chromatography (SEC) for Nup145C-Sec13$_{fusion}$+Nup84$_{1-424}$ (blue), pre-incubated with VHH-SAN8 (1:2 molar ratio) (red), or pre-incubated with VHH-SAN8 and VHH-SAN9 (1:1:1 molar ratio) (green). (b) SDS-PAGE analysis of the SEC fractions indicated for each SEC experiment. *indicates contaminant protein. Co-elution of VHH-SAN8 and VHH-SAN8/9 shows that both nanobodies can bind Nup84 while in complex with Nup145C.

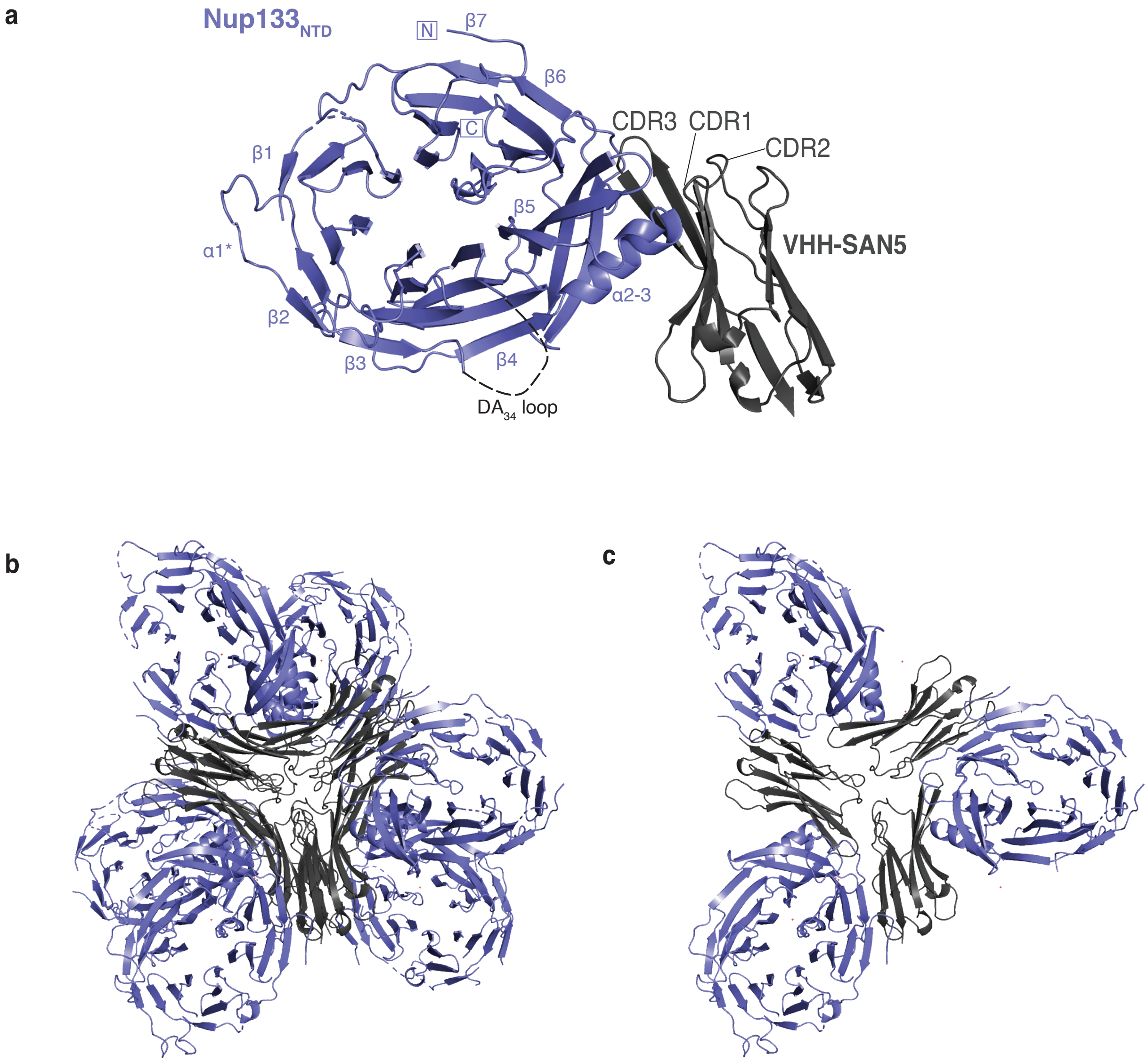


### Supplementary Figure 7.

**Structure of Nup133$_{NTD}$-VHH-SAN5 complex.**

(a) Nup133$_{NTD}$-VHH-SAN5, with Nup133$_{NTD}$ in purple and VHH-SAN5 in dark gray. N and C termini are indicated, and β-sheets and α-helices are labeled on Nup133$_{NTD}$. Complementarity determining region (CDR) loops are labeled on VHH-SAN5. (b) Packing of Nup133$_{NTD}$-VHH-SAN5 in the asymmetric unit. Colors as in (a). (c) Heterohexameric assembly within the asymmetric unit highlighting packing interfaces mediated by VHH-SAN5.

### Supplementary Table 1. BackPhyre Structural Homology Results

Table includes hits that cover >100 amino acids and >30% confidence.

| Protein | Description | Alignment coverage (%) | Confidence | % Sequence ID |
|---|---|---|---|---|
| Nic96 | Scaffold nucleoporin, Nic96/inner ring complex | 224-637 (63) | 98.7 | 18 |
| Nup145C | Scaffold nucleoporin, Y complex | 1,027-1,284 (39) | 97.2 | 16 |
| Sec31 | COPII vesicle coat component | 643-875 (35) | 95.1 | 18 |
| Sea4 | SEA complex, lysosomal membrane interaction and autophagy | 772-996 (34) | 79.0 | 25 |
| Sec16 | COPII vesicle coat assembly | 255-340 (13) | 72.5 | 17 |
| Nup85 | Scaffold nucleoporin, Y complex | 465-566 (15) | 31.5 | 14 |
